# Phylostems: a new graphical tool to investigate temporal signal of heterochronous sequences at various evolutionary scales

**DOI:** 10.1101/2020.10.19.346429

**Authors:** Anna Doizy, Amaury Prin, Guillaume Cornu, Frederic Chiroleu, Adrien Rieux

## Abstract

1. Molecular tip-dating of phylogenetic trees is a growing discipline that uses DNA sequences sampled at different points in time to co-estimate the timing of evolutionary events with rates of molecular evolution. Such inferences should only be performed when there is sufficient temporal signal within the analysed dataset. Hence, it is important for researchers to be able to test their dataset for the amount and consistency of temporal signal prior to any tip-dating inference. For this purpose, the most popular method considered to-date has been the “root-to-tip regression” which consist in fitting a linear regression of the number of substitutions accumulated from the root to the tips of a phylogenetic tree as a function of sampling times. The main limitation of the regression method, in its current implementation, relies in the fact that the temporal signal can only be tested at the whole-tree evolutionary scale.
2. To fill this methodological gap, we introduce phylostems, a new graphical and user-friendly tool developed to investigate temporal signal at every evolutionary scale of a phylogenetic tree.
3. Phylostems allows detecting without *a priori* whether any subset of a tree would contain sufficient temporal signal for tip-based inference to be performed. We provide a “how to” guide by running phylostems on empirical datasets and supply guidance for results interpretation. Phylostems is freely available at https://pvbmt-apps.cirad.fr/apps/phylostems.
4. Considering the impressive increase in availability and use of heterochronous datasets, we hope the new functionality provided by phylostems will help biologists to perform thorough tip-dating inferences.

## Introduction

“Tip-dating” of phylogenetic trees is a popular and powerful type of genetic analysis aiming to make use of sequence data isolated at different points in time (i.e., heterochronous datasets) to co-estimate the timing of evolutionary events with rates of molecular evolution (Rieux & Balloux, 2016). Tip-dating requires working on measurably evolving populations (MEPs) which consist in datasets displaying detectable amounts of *de novo* nucleotide changes among the DNA sequences sampled at different timepoints (Drummond, Pybus, Rambaut, Forrest, & Rodrigo, 2003). Our ability to capture measurable amount of evolutionary change from sequence data is a factor of various parameters including the evolutionary rate per site per unit of time (*μ*), the width of the sampling interval (*t*), the number of sites in the sequences (*L*) and the time to the Most Recent Common Ancestor (MRCA) of all sequences (*T_MRCA_*). Originally, only fast-evolving organisms such as RNA viruses were classifiable as MEPs but the recent rise in our ability to sequence DNA at high throughput from both modern and ancient material has led to a massive increase in both sequence length (*L*) and the timespan covered by the sequences (*t*), hence opening up the field of tip-dating to a variety of additional organisms (Biek, Pybus, Lloyd-Smith, & Didelot, 2015).

Phylogenetic inferences performed on such time-structured sequence data represent a powerful tool for hypothesis testing (Rieux & Balloux, 2016). They have notably been critical for *i)* dating key events in human evolutionary history (Fu et al., 2013; Rieux et al., 2014), *ii)* improving our understanding of various important pathogens emergence, spread and evolution (Bos et al., 2014; Faria et al., 2014; Eldholm et al., 2015; O’Hanlon et al., 2018; Vanhove et al., 2019; Rambaut, 2020), *iii)* investigating the relative impacts of climatic and anthropogenic factors on the widespread extinctions of large mammals (Shapiro et al., 2004; Stiller et al., 2010), *iv)* providing meaningful information about pathogens host species jumps (Weinert et al., 2012) and *v)* estimating unknown sequence’s ages in various organisms (Shapiro et al., 2011).

Inferences from tip-calibrated phylogenetic trees should only be performed when there is sufficient temporal signal within the analysed dataset (Drummond, Pybus, Rambaut, et al., 2003; Duchêne, Duchêne, Holmes, & Ho, 2015; Murray et al., 2016; Rieux & Balloux, 2016). This will for instance not be the case if the sampling period is too short for sufficient evolutionary changes to be measured, if evolutionary rates are too low or variable amongst lineages or if some samples have incorrectly been dated (Rambaut, Lam, Carvalho, & Pybus, 2016). As such it is important for researchers to be able to test their dataset for the amount and consistency of temporal signal prior to any tip-dating inference. For this purpose, the most popular method considered to-date has been the “root-to-tip regression” which consist in fitting a linear regression of the number of substitutions accumulated from the root to the tips of a phylogenetic tree as a function of sampling times (Buonagurio et al., 1986; Shankarappa et al., 1999; Korber et al., 2000; Drummond, Pybus, & Rambaut, 2003). If sampling dates are sufficiently different, then more recently sampled sequences should have undergone substantially more evolutionary change than earlier sampled sequences, which would result in a positive correlation. This method has often been used as a diagnostic of data quality and of the reliability rate estimates, where the slope coefficient corresponds to the substitution rate under the assumption of a strict molecular clock, the X-intercept is an estimate of the date of the root of the tree and R^2^ indicates the degree to which sequence evolution has been clocklike. However, the root-to-tip regression method is not statistically suitable for proper hypothesis testing because the individual data points are not independently distributed, and are instead partially correlated due to their phylogenetic shared ancestry (Drummond, Pybus, & Rambaut, 2003). To overcome this limitation, Navascues et al. (2010) suggested a non-parametric approach using permutations to test whether the correlation is stronger than expected if the sampling dates were randomly assigned. More recently, other phylogenetic approaches such as the date-randomization test (Ramsden, Melo, Figueiredo, Holmes, & Zanotto, 2008; Duffy & Holmes, 2009; Duchêne et al., 2015; Murray et al., 2016) or model selection/comparison (Rambaut, 2000; Murray et al., 2016; Duchene et al., 2019), although way more computationally intensive, have also been introduced and shown to be more robust tests for temporal signal detection and characterization.

Despite its statistical pitfalls, the regression method remains a very helpful exploration tool to quickly assess the extent of temporal signal within a dataset. It only requires a rooted molecular phylogeny (whose branch lengths represent genetic distance) estimated from heterochronous (dated) sequences and runs instantaneously. The regression method has been implemented in the popular and interactive graphical program TempEst (Rambaut et al., 2016), formerly known as Path-O-Gen. The main limitation of the regression method in its current implementation relies in the fact that the temporal signal can only be tested at the whole-dataset (tree) evolutionary scale. However, although a significant positive correlation would indicate the presence of detectable amounts of *de novo* mutations within a tree timescale, a non-positive (or a statistically non-significant) correlation does not necessarily mean that no temporal signal exists at a reduced timescale, as illustrated in Fig 1. To fill this methodological gap, we introduce phylostems, a new graphical and user-friendly tool developed to investigate temporal signal at every evolutionary scales of a phylogenetic tree. Phylostems allows detecting without *a priori* whether any subset of a tree would contain sufficient temporal signal for tip-based inference to be performed. We provide a “how to” guide by running phylostems on empirical datasets and supply insights on interpreting the outputs.

**Fig. 1.**
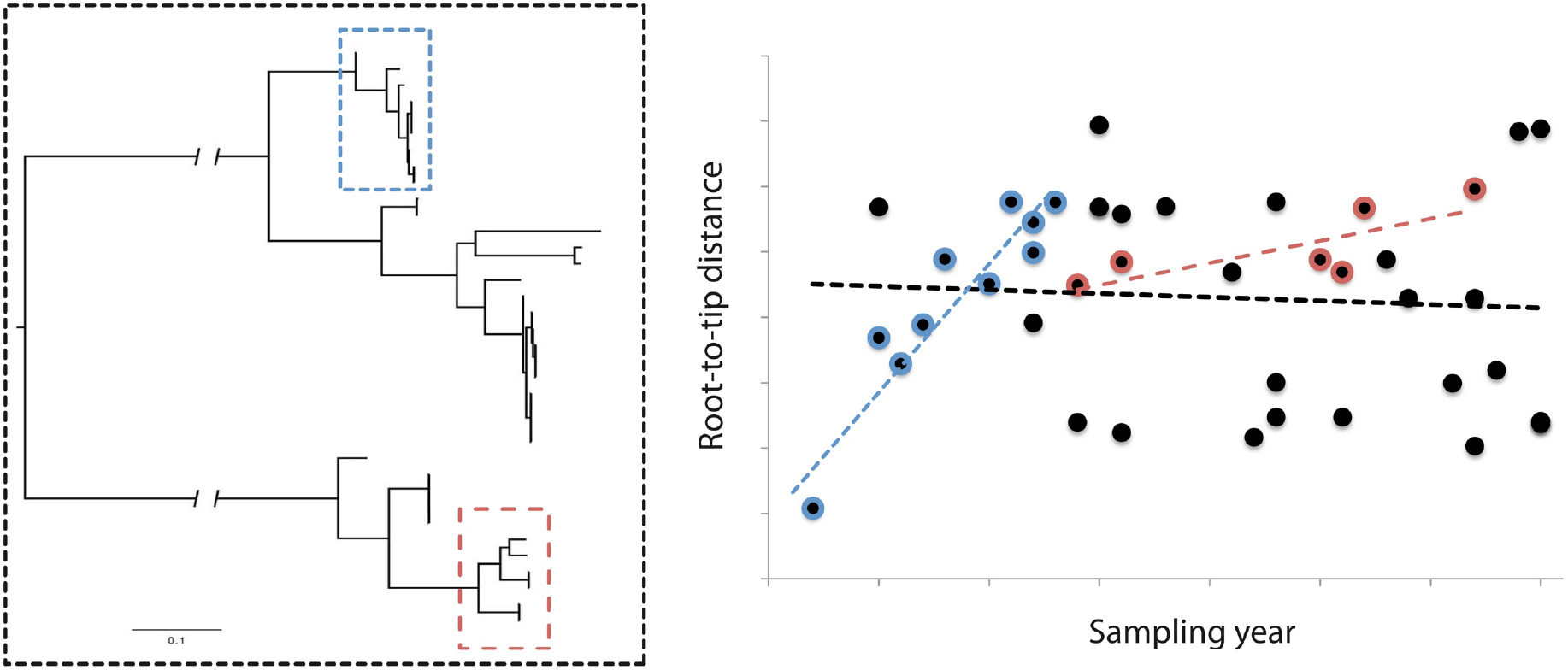
Let’s a tree (left panel) be constructed from a dataset of heterochronous sequences. When investigating temporal signal on the whole dataset using the regular root-to-tip regression method (right panel), no significant signal was found as the slope of the regression (black dotted line) appears to be non-positive. Hence, tip-based inferences should not be performed at the whole dataset timescale. However, as illustrated by the red and blue positive regression slopes calculated on two subsets of samples (red and blue squares on the tree), positive temporal signal exists at reduced evolutionary timescales at which thorough tip-based inferences could be performed. The main objective of phylostems is to provide the user with a graphical tool to detect without *a priori* such evolutionary clades.

## Materials and Methods

The program phylostems (Phylogenetic Scaling of Temporal Signal) is an open source, graphical Shiny based R application (Chang, Cheng, Allaire, Xie, & McPherson, 2018; R Core Development Team, 2020) built for exploring temporal signal at various scales of a phylogenetic tree. Shiny is an R package that makes it easy to build interactive web applications from R (https://shiny.rstudio.com/). Phylostems can be either used online at https://pvbmt-apps.cirad.fr/apps/phylostems/ or executed locally by downloading its source code from https://gitlab.com/cirad-apps/phylostems. A schematic representation of phylostems workflow is presented in Fig. 2.

**Fig. 2.**
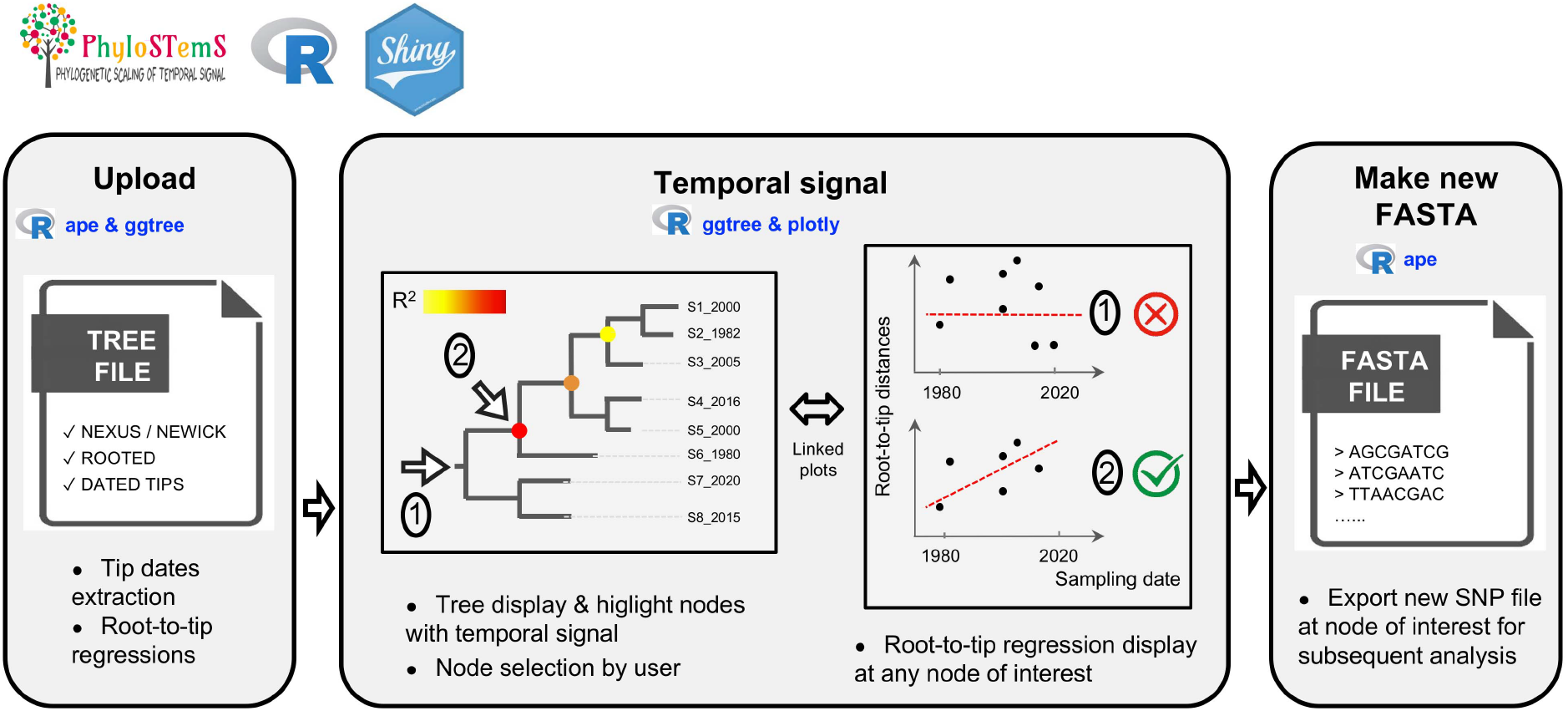
Schematic representation of phylostems workflow. Main boxes (“Upload”, “Temporal signal” & “Make new FASTA”) represent the internal structure of the application organized in three main panels. Major tasks performed in each panel are summarized along with sourced R packages.

As input, phylostems requires *(i)* a phylogenetic tree in computer-readable Nexus or Newick format with branch lengths scaled as genetic distances only, such as the ones computed using maximum likelihood approaches (e.g. Guindon et al., 2010; Minh et al., 2020; Stamatakis, 2014). In its current implementation, the online version of phylostems allows uploading trees with 1500 sequences at maximum. Larger trees will need to be processed locally by sourcing the gitlab version. *(ii)* Prior to be loaded in phylostems, the tree needs to be rooted, either at a position chosen by the user (with an outgroup) or at a most compatible location with the assumption of a strict molecular clock (using for instance the rtt function from the ape R package (Paradis & Schliep, 2019)). When possible, we advise to use outgroup-rooted trees. Finally, *(iii)* sampling/isolation dates needs to be known for each sequences and specified within tip labels. Before-Christ (B.C) dates, sometimes required to handle sequences generated from ancient DNA data can be specified using negative values (e.g. - 400.5). Note that since missing dates are not allowed, sequences with unknown sampling years needs to be pruned out from the tree (using for instance the drop.tip function from the ape R package) prior to be uploaded in phylostems.

When a tree has correctly been loaded in phylostems, a distribution of sampling dates is plotted within the “upload” panel allowing for a visual check of sequences temporal width. At this stage, the phylogenetic tree has been loaded using the ape R package and root-to-tip distances for all sequences are recorded. Temporal signal is hence tested at every node of the input tree (including its root) meeting the following conditions required to perform a linear regression: *i)* the node must be the parent of at least n=3 tips, *ii)* there should be at least n=3 distinct combination of root-to-tip distances and sampling dates and *iii)* there should be at least n=2 different sampling dates. At each of such nodes, linear regression between sampling dates and root-to-tip distances is performed and the following parameters: (1) p-value, (2) slope, (3) adjusted R^2^, and (4) intercept with the x-axis values are recorded.

Phylostems’s main results are provided within the “Temporal signal” panel. First an annotated phylogenetic tree is interactively plotted by sourcing both ggtree and plotly packages (Yu, Smith, Zhu, Guan, & Lam, 2017; Sievert, 2020). On this tree, nodes with temporal signal, *i.e.* nodes at which root-to-tip linear regression yielded a statistically significant and positive slope, are highlighted with colours scaling to R^2^ value. The default threshold for the linear regression p-value has been fixed to 0.05 but the user can interactively modify it using a slider bar, which enable easy investigation of nodes with borderline significant trends. A table summarizing the nodes with temporal signal is also displayed along with respective number of descending sequences, p-value, slope and adjusted R^2^ values. Most importantly, phylostems allow the user to visualize the root-to-tip regressions at any chosen node of interest. To do so, the user simply needs to click on a node, and the associated root-to-tip regression will be displayed. Both the tree and the root-to-tip regression plots are linked, so that data points (or tree tips) selected in one plot will automatically be highlighted on the other one. This enables easy investigation of outliers and sequences or clades of interest.

Finally, when temporal signal is found at the within-tree scale, phylostems’s “Make new FASTA” panel allows generating a new subset sequence FASTA file that only include the variant sites for the descending tips of a node of interest, a dataset suitable for further tip-dating inferences.

In the following, we use two previously published empirical datasets to illustrate how phylostems allows users exploring temporal signal at various evolutionary scales within phylogenetic trees. For both datasets, we downloaded rooted-ML tree files built from non-recombining genomic sequences from their original publications. The first dataset contains 45 strains of *Xyllela fastidiosa* (hereafter *Xf*) sampled worldwide between 1983 and 2016 (Vanhove et al., 2019). *Xf* is a bacterial crop pathogen of global importance, currently threatening agriculture in various Europeans countries (Sicard et al., 2018). The second dataset comprises 98 hantaviruses isolates sampled from bank voles in Belgium between 1984 and 2016 (Laenen et al., 2019). Hantaviruses are important zoonotic viral pathogens that can cause hemorrhagic fever with renal syndrome and pulmonary syndrome, potentially life-threatening diseases in humans (Maes, Clement, Gavrilovskaya, & Van Ranst, 2004).

## Results

### *Xf* dataset

We first loaded the *Xf* rooted tree within phylostems ‘s “upload” panel (see Fig. 3). Looking at the plot of the sampling dates distribution, one can perform a quick visual check of sequences temporal width (here 1983-2016) to validate the data importation process. Moving to the “Temporal signal” panel, phylostems displays the *Xf* phylogenetic tree on which the structuration by the four subspecies: ssp. *pauca*, *multiplex*, *morus*, *fastidiosa* can be easily distinguished (see Fig. 4A). Visual inspection of the *Xf* tree in phylostems demonstrated a lack of strong and deep temporal signal as neither the root nor the MRCA of each subspecies displayed any significant correlation between root-to-tip distances and sampling ages, as highlighted by the absence of annotations at those nodes. Phylostems detected only one internal node (node 82) associated with temporal signal within the *Xf* tree. This node is the MRCA of a small clade containing 9 samples within the *Xf pauca ssp* clade. When clicking on this node, phylostems displays the associated root-to-tip regression plot and parameters (R^2^ = 0.38, slope = 6.9E-7, P-val = 0.045, see Fig.4 B). According to phylostems’s results, this small clade (N=9 sample) is the only evolutionary scale suitable for phylogenetic tip-based inferences in BEAST or other programs within the *Xf* dataset. To do so, the “Make new FASTA” panel allows generating a new sequence file that only include the variant sites for the 9 *Xf* samples within the clade with detected temporal signal.

**Fig. 3.**
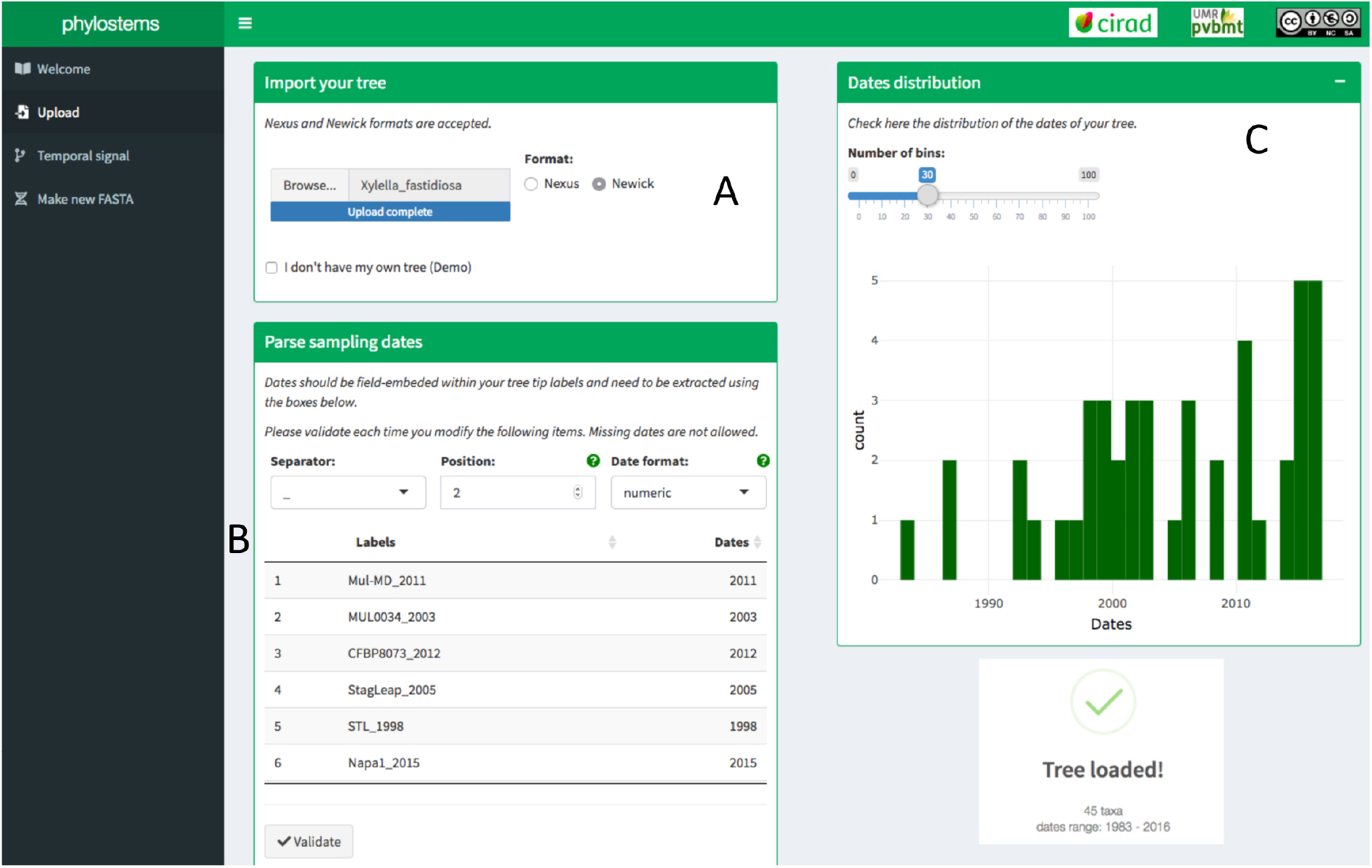
Phylostems ‘s upload panel requesting the user to load a phylogenetic tree (A) and specify tip sampling dates from field-embedded values (B). Once loaded, a distribution of sampling dates is plotted allowing for a visual check of sequences temporal width (C).

**Fig. 4.**
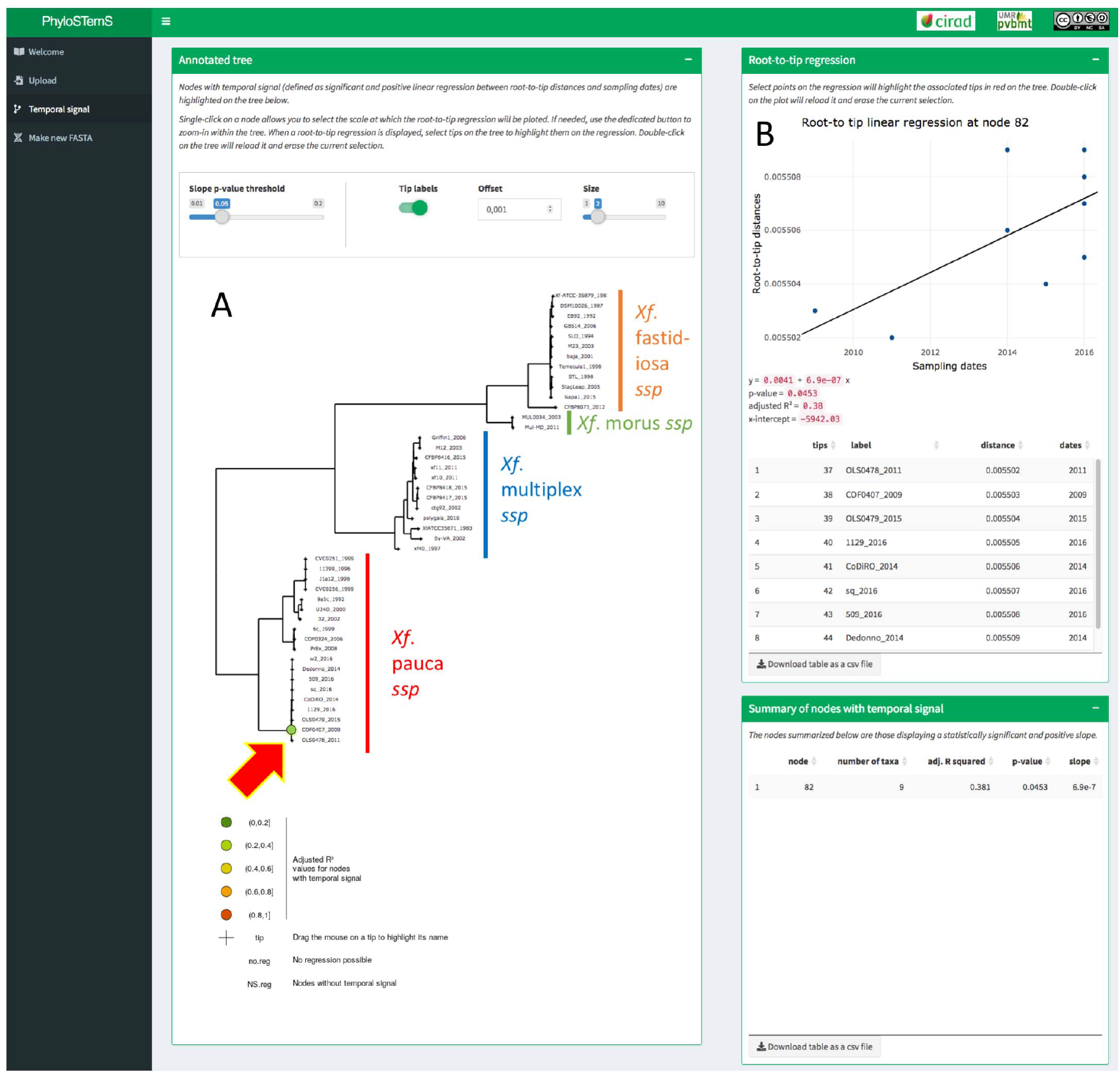
Annotated phylogenetic tree of *Xylella fastidiosa* empirical dataset (A). Red arrow indicates the only node at which temporal signal was found. Root-to-tip regression at this node, along with associated parameters are plotted in (B).

### Hantaviruses dataset

Visual inspection of the Hantaviruses tree in phylostems demonstrated heterogenous temporal signal amongst clades, here referring to three geographical sampling areas namely Ardennes, Campine and Sonian Forest (Fig 5.A). Phylostems revealed a lack of temporal signal both at the whole tree scale and for the Sonian Forest clade. Temporal signal was observed at the MRCA of the Campine and Ardennes clades as well as within the Ardennes clade, as represented by the several highlighted nodes on the tree. A table listing all the nodes associated with temporal signal along with their associated statistics is given in Fig 5.B. When plotting the regression at the MRCA of the Campine and Ardennes clades, phylostems allows visually identifying outlier samples that are significantly deviating from the root-to-tip regression line (Fig 5.C). Here, all outliers felt within the Campine clade, suggesting that phylogenetic tip-based inferences should not be performed on both the Campine and Ardennes clades simultaneously. Possible causes for such outliers are multiple and will be argued in the discussion section.

**Fig. 5.**
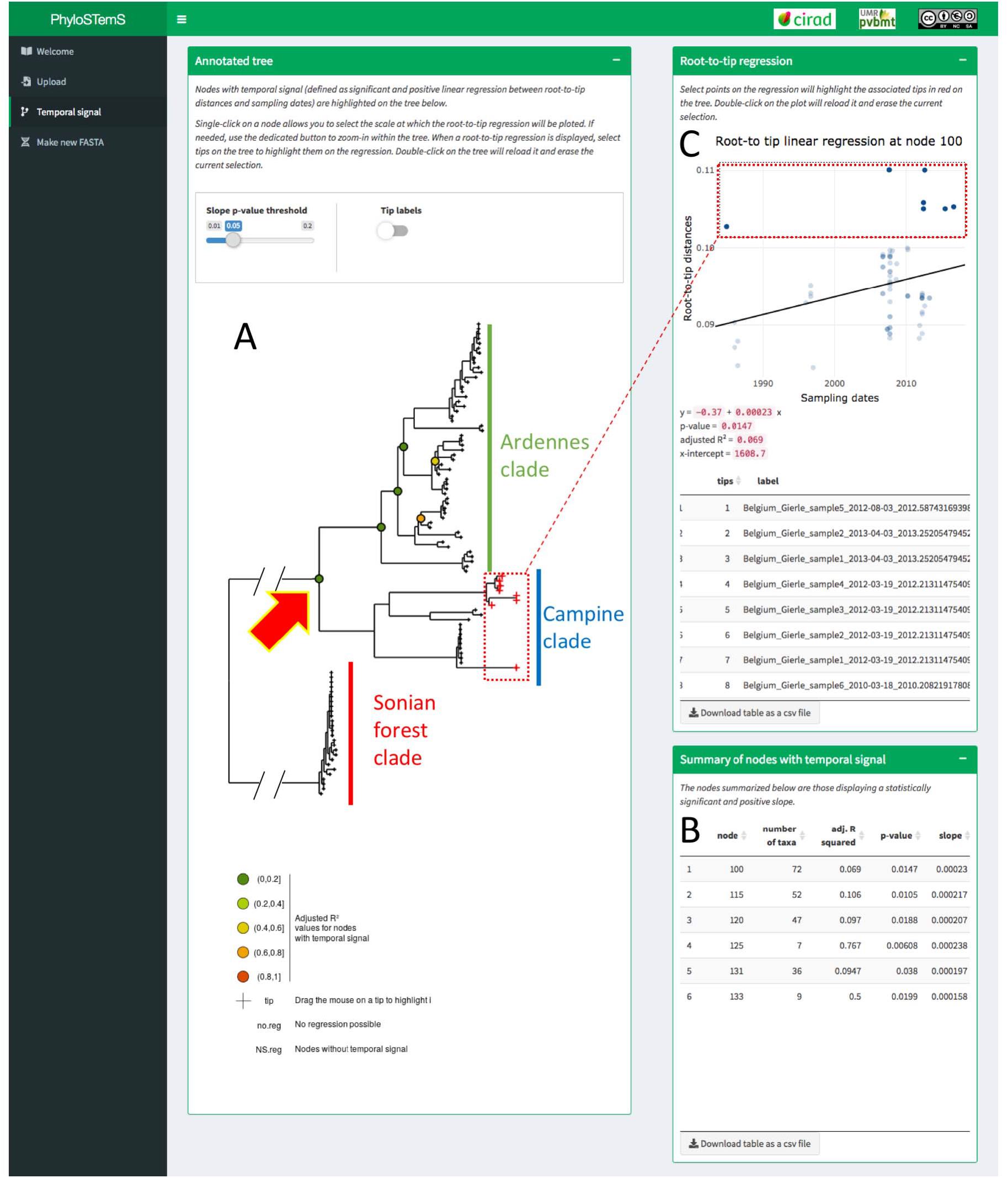
Annotated phylogenetic tree for the Hantaviruses empirical dataset (A). Coloured circles indicate nodes at which temporal signal was found. A table summarizing those nodes, along with associated linear regression parameters is given in (B). Root-to-tip regression at node highlighted by the red arrow is plotted in (C). Both the tree and the regression plots are linked, so that data points (or tree tips) selected in one plot will automatically be highlighted on the other one, as illustrated by the red-dotted frames.

## Discussion

We introduce phylostems, a new graphical and user-friendly tool developed to investigate temporal signal within phylogenetic trees using the root-to-tip regression method. Previous implementations of this method, such as for instance in the popular and interactive graphical program TempEst (Rambaut et al., 2016) were designed to test temporal signal at the whole tree scale (i.e. at its root). Investigating temporal signal at smaller evolutionary scales was previously doable, but this task required the user to *i) a priori* decide at which scale (i.e. on which samples) performing the test and *ii)* manually splitting or reconstructing the tree for every of such scales. The main improvement of phylostems is to allow detecting, in a single step and without *a priori*, any evolutionary scale at which temporal signal may exist within a phylogenetic tree.

Exploring the degree of temporal signal in heterochronous sequences datasets before proceeding to inference using formal molecular clock models is a crucial task (Rieux & Balloux, 2016). As illustrated by the two empirical datasets analyzed in this study, temporal signal may sometimes be heterogeneous within a tree with substantial differences between clades. In such cases, we hope that phylostems will help researchers detecting the most appropriate scales, if any, at which thorough tip-based inferences may be performed. However, because of the statistical pitfalls associated with the root-to-tip regression method (Rambaut, 2000; Rambaut et al., 2016), phylostems should rather be seen as a fast, visual and qualitative data exploration tool for temporal signal detection but should not be used to test hypotheses or undertake statistical model selection. Once temporal signal has been detected in phylostems, we advise users to make use of other available methods such as non-parametric permutations (Navascués et al., 2010), date-randomization test (Ramsden et al., 2008; Duffy & Holmes, 2009; Duchêne et al., 2015; Murray et al., 2016) or model selection/comparison (Murray et al., 2016; Duchene et al., 2019) to validate the existence of measurably evolving populations in their datasets.

Finally, phylostems can also help identifying outliers or groups of samples that substantially differ from the root-to-tip regression line and may require careful handling to avoid bias during phylogenetic inferences. First, as illustrated by the analyse of the Hantaviruses dataset, different clades or populations in a tree may be characterized by positive but contrasted root-to-tip regression patterns that might arise from sampling bias or differences in life-history traits between clades (e.g. environmental factors, population density, evolutionary rates or epidemiological parameters). In such a case, it is suggested to perform independent phylogenetic inferences on each clade/population (Laenen et al., 2019). In other cases, outlier sequences whose sampling date is incongruent with their genetic divergence and phylogenetic position can be spotted from the regression plot (Rambaut et al., 2016). Such anomalies can reflect a problem with *i)* the sequence itself (e.g. low quality, sequencing/assembly/alignment errors, recombination or hypermutation) or *ii)* the sampling date(s) (e.g. mislabelling or biological contamination). Should the case of such outlier sequences arise, those samples should be excluded from subsequent phylogenetic inferences.

Considering the impressive increase in availability and use of heterochronous datasets, we hope the functionality provided by phylostems will help users to perform thorough tip-dating inferences. Pylostems is a dynamic application by nature. New functions will be added as new needs arise.

## Acknowledgements

This work was financially supported by l’Agence Nationale pour la Recherche (JCJC MUSEOBACT contrat ANR-17-CE35-0009-01), the European Regional Development Fund (ERDF contract GURDT I2016-1731-0006632), Région Réunion and the French Agropolis Foundation (Labex Agro – Montpellier, E-SPACE project number 1504-004). We are grateful to S. Falala for his advices on building Shiny apps and CIRAD for providing hosting of the application server. We thank R. Almeida, M. Vanhove & B. Vrancken for providing access to the empirical datasets analyzed in this study. We also thank L. van Dorp, P. Campos, C.G. Crego, E. Conte, T.T CAO, F. Balloux & D. Richard for interesting discussions and testing previous versions of the app on their own datasets.

## Data availability

Phylostems can be executed online at https://pvbmt-apps.cirad.fr/apps/phylostems/ but source code can also be downloaded from https://gitlab.com/cirad-apps/phylostems for local implementation. The two empirical trees used in this paper (Hantaviruses and *Xylella fastidiosa*) are accessible from the gitlab repository.

## Authors contribution

A.R initially conceptualized the method. A.P generated a first version of the code. A.D improved it and converted it into a Shiny application with advices from A.R, G.C & F.C. G.C managed the online implementation & maintenance of the app. A.D & A.R wrote the first draft and all authors contributed to the final version.

## Notes

https://pvbmt-apps.cirad.fr/apps/phylostems

### Competing Interest Statement

The authors have declared no competing interest.

